# A comparative analysis reveals irreproducibility in searches of scientific literature

**DOI:** 10.1101/2020.03.20.997783

**Authors:** Gábor Pozsgai, Gábor L. Lövei, Liette Vasseur, Geoff Gurr, Péter Batáry, János Korponai, Nick A. Littlewood, Jian Liu, Arnold Móra, John Obrycki, Olivia Reynolds, Jenni A. Stockan, Heather VanVolkenburg, Jie Zhang, Wenwu Zhou, Minsheng You

**Affiliations:** State Key Laboratory of Ecological Pest Control for Fujian and Taiwan Crops, Institute of Applied Ecology, Fujian Agriculture and Forestry University, Fuzhou 350002, China; Joint International Research Laboratory of Ecological Pest Control, Ministry of Education, Fuzhou 350002, China; Department of Agroecology, Flakkebjerg Research Centre, Aarhus University, 4200 Slagelse, Denmark; UNESCO Chair on Community Sustainability: From Local to Global, Dept. Biol. Sci., Brock University, Canada; Graham Centre for Agricultural Innovation (Charles Sturt University and NSW Department of Primary Industries), Orange NSW 2800, Australia; Agroecology, University of Goettingen, 37077 Goettingen, Germany; “Lendület” Landscape and Conservation Ecology, Institute of Ecology and Botany, MTA Centre for Ecological Research, 2163 Vácrátót, Hungary; Department of Water Supply and Sewerage, Faculty of Water Science, National University of Public Service, 6500 Baja, Hungary; Department of Environmental Sciences, Sapientia Hungarian University of Transylvania, 400193 Cluj-Napoca, Romania; Eötvös Lórand University, 1053 Budapest, Hungary; University of Cambridge, Cambridge, UK; University of Pécs, 7622 Pécs, Hungary; University of Kentucky, Lexington, USA; cesar, 293 Royal Parade, Parkville, Victoria 3052, Australia; Biosecurity and Food Safety, NSW Department of Primary Industries, Narellan, NSW 2567, Australia; Ecological Sciences, The James Hutton Institute, Aberdeen, UK; State Key Laboratory of Rice Biology, Key Laboratory of Molecular Biology of Crop Pathogens and Insects, Ministry of Agriculture, Zhejiang University, Hangzhou, China

**Keywords:** Database, search engine, search location, repeatability

## Abstract

Repeatability is the cornerstone of science and it is particularly important for systematic reviews. However, little is known on how database and search engine choices influence replicability. Here, we present a comparative analysis of time-synchronized searches at different locations in the world, revealing a large variation among the hits obtained within each of the several search terms using different search engines. We found that PubMed and Scopus returned geographically consistent results to identical search strings, Google Scholar and Web of Science varied substantially both in the number of returned hits and in the list of individual articles depending on the search location and computing environment. To maintain scientific integrity and consistency, especially in systematic reviews, action is needed from both the scientific community and scientific search platforms to increase search consistency. Researchers are encouraged to report the search location, and database providers should make search algorithms transparent and revise access rules to titles behind paywalls.

## Introduction

Since the 17^th^ century and Newton’s strict approach to scientific inquiry[1], research has increasingly relied on rigorous methodological constrains. One of the cornerstones of the scientific method is reproducibility. However, a recent study shows that most scientists believe that a substantial proportion of methods published in peer-reviewed papers are not reproducible, creating a ‘reproducibility crisis’[2]. Following similar arguments, narrative reviews are increasingly being replaced by systematic reviews, also called “evidence-based synthesis”[3]. Transparency and repeatability are also cornerstones of this method of knowledge synthesis. However, the repeatability of systematic reviews remains rarely examined. Though repeatability in such studies is of utmost importance, and detailed protocols are available[4,5], the technical aspects of these underpinning databases and search engines have not been systematically tested and, at present, there is no recommendation on these technical aspects.

As primary scientific literature is rapidly expanding[6], scientists are unable to keep track of new discoveries by focusing only on the primary literature[7,8], so systematic reviews have become increasingly important[9]. Recognized weaknesses of the traditional, narrative reviews include the non-transparency of the literature selection process, evaluation criteria, and eventual level of detail devoted to individual studies[10]. With the advent and rapid development of Internet-based databases and search engines, the role of narrative reviews is now being overtaken by new, quantitative methods of evidence synthesis[11,12]. A core requirement in these activities, repeatability, crucially depends on reliable databases[13]. Large scientific databases/search engines, such as PubMed, Web of Science and Scopus, are essential in this process. They have been primary electronic search engines for scientists since 1997 with the inauguration of PubMed[14]. Today, nearly all scientists working on various forms of evidence-based synthesis use these databases/search engines to find relevant papers as the basis for further analysis.

An important condition in the whole process is that the evidence base must be solid: a given search string in a database should generate identical results, independent of search locations, provided the searches are running at the same time. If this assumption were violated, it would have serious consequences for the reliability and repeatability of the data and papers selected for a specific systematic review. Therefore, there is a need to know what variables and/or parameters should be included in the methodology of any search to ensure its repeatability. One of the most crucial steps is to define which database and engine search is going to be used for obtaining the data to be synthesized.

Differences among the most commonly used scientific search engines and databases are well documented[13,15,16] but knowledge of the consistency within databases in relation to geographical location where the search is requested from (but see Gusenbauer and Haddaway[13]), software environment, or computer configuration remain surprisingly limited. Since the search histories of users may be stored in the browsers’ cache, and considered by the scientific search engines, repeated and identical searches may result in different outcomes. During a recent systematic review in ecology, we accidentally discovered that a multi-locus search performed on 1 February 2018, using an identical search string in Web of Science, produced radically different number of hits at different institutions at Hangzhou and Fuzhou, in China, and in Denmark (2,394, 1,571, and 7,447, respectively).

Since there is no known study comparing the consistency of returned papers over successive identical searches using several databases in one machine, we examined the way search engines deliver results and decided to systematically explore the inconsistencies found. Our study aimed to evaluate the consistency of search engines by comparing the outcomes from identical search strings ran on different computers from a wide range of localities across the world, with various software backgrounds, and using different search engines.

To investigate the repeatability of scientific searches in four of the major databases and search engines, Web of Science, Scopus, PubMed, and Google Scholar, we generated search strings with two complexity levels in ecology and medicine and ran standardized searches from various locations in the world, within a limited timeframe. According to our null hypothesis, every search engine should give the exact same number of results to the same search (after the search term has been adjusted to match the specific requirements for each of these search engines), and therefore, a metric, showing the proportional deviance of the search hits, should always be zero. We, therefore, first tested if summarized *average absolute deviation proportions* (AADPs) for each search engine were significantly different from the ideal value (zero) by using robust non-parametric tests. AADPs of search engines were compared to each other and factors driving the differences were investigated. Similarly, the publications found by any given search engine from identical searches should be also identical, thus, the mean similarities between search runs should be 100%, and the scatter of the ordinated points should be zero. In order to test whether these requirements were met, Jaccard distances[17] of the first twenty hits were used for within and between group ordinations and multivariate analysis.

## Results

Our time-synchronized, cross-institution and multi-location search exercise resulted in a large variation among the hits obtained using any of the search terms. Google Scholar generally yielded a greater number of hits than any other databases for all the locations (Table 1). As expected, less complex and medical search terms tended to result in greater hit numbers than complex ecological ones.

**Table 1.**
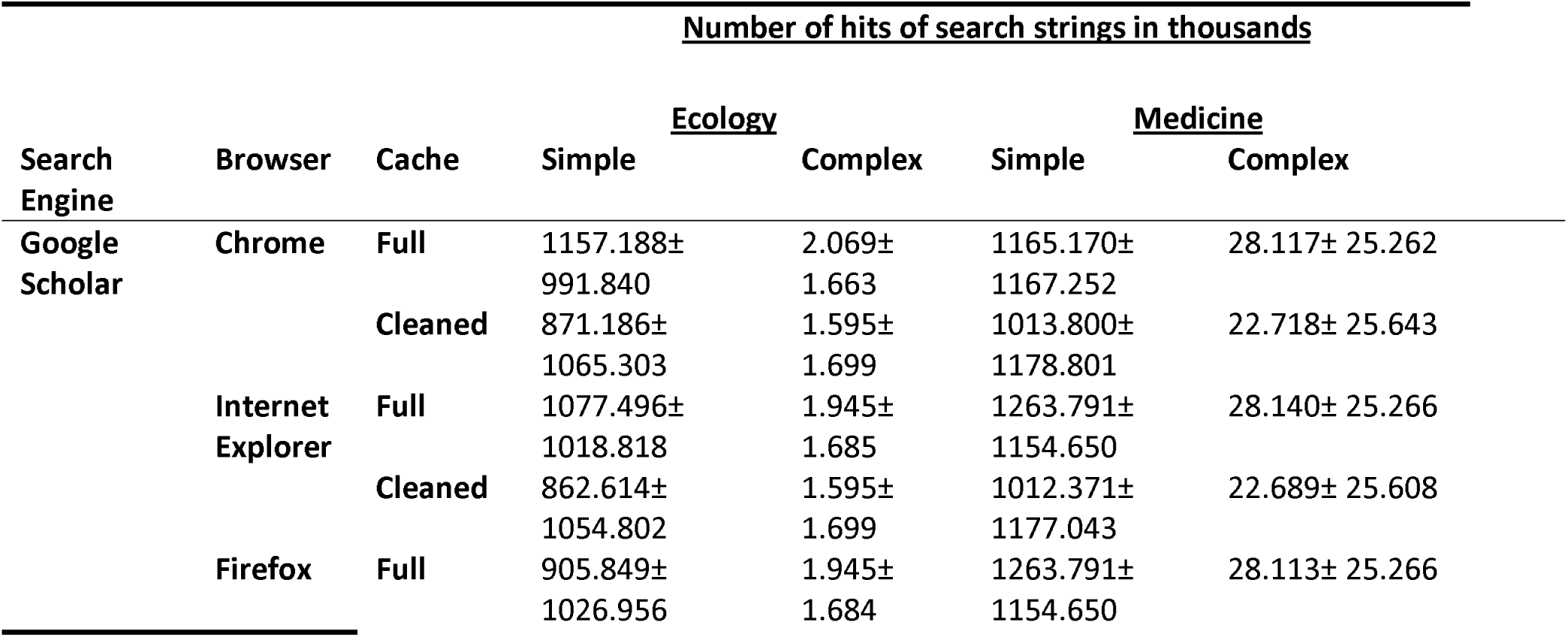
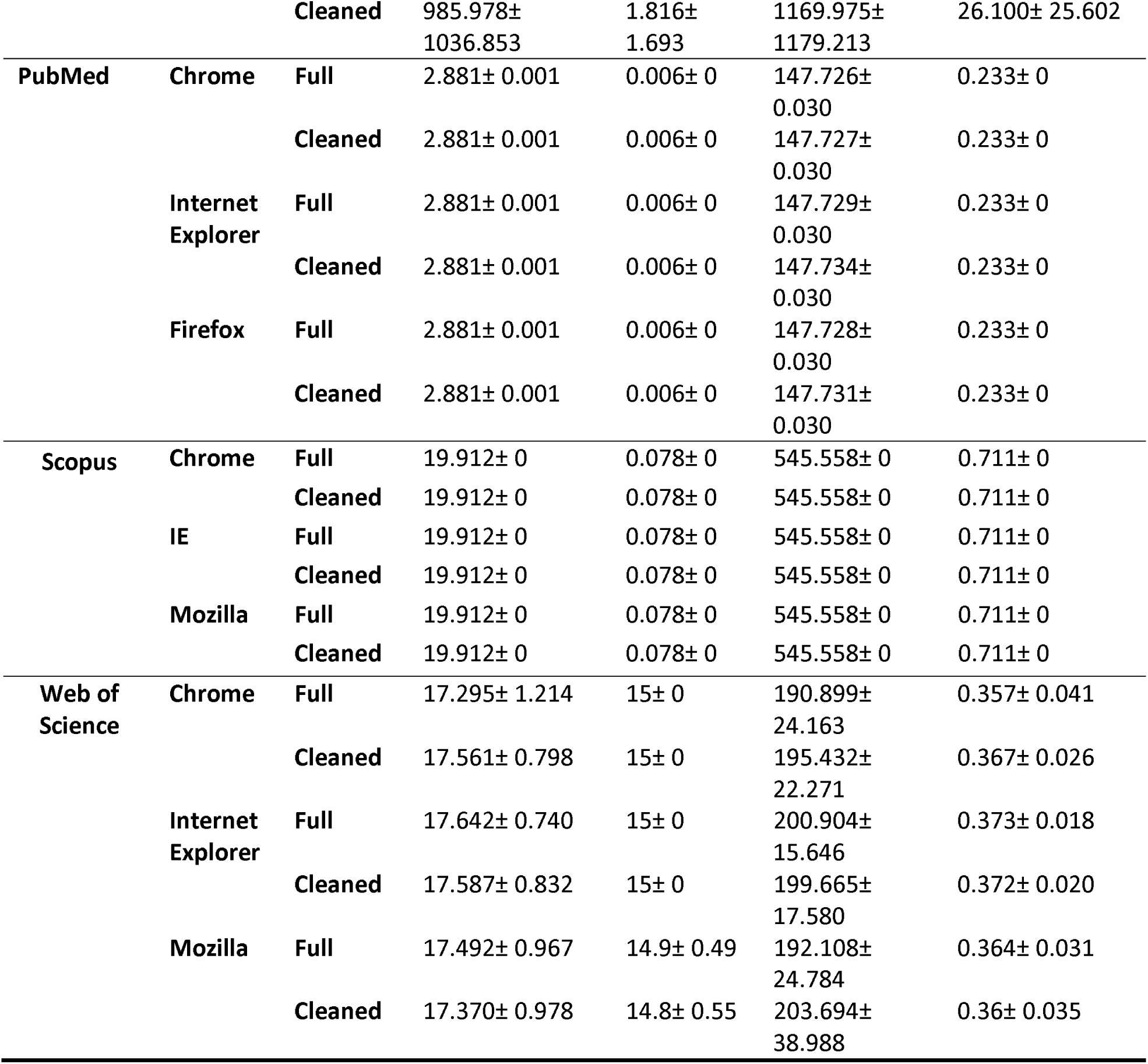
Comparison of the mean numbers of hits (SD) resulting from simple vs. complex search strings in the fields of ecology and medicine using different search engines, different browsers and cache handling

The AADP (see Materials and Methods) of every search engine and database, except Scopus, significantly deviated from the desirable zero (Table 2). However, we have noticed that both PubMed and Web of Science were updated during the search process, at 17:00 GMT and 19:00 GMT, respectively. When the results from PubMed and Web of Science were split into two groups, before and after the time of the daily update, none of the AADPs from PubMed searches significantly differed from zero. In contrast, the results from Web of Science searches consistently showed significant deviation, indicating inconsistency in the number of returned hits by search location.

**Table 2.** Mean and standard deviations of recorded average absolute deviation proportions (AADP) for each investigated search engines, separated by search topic and search expression complexity. Values are shown in percentage.

| Topic/Complexity | GScholar | PubMed | Scopus | WoS |
| --- | --- | --- | --- | --- |
| Ecology/Complex | 85.319±9.426 | 0.000±0.000 | 0.000±0.000 | 0.629±1.964 |
| Ecology/Simple | 98.107±4.063 | 0.035±0.000 | 0.000±0.000 | 4.009±3.459 |
| Medicine/Complex | 90.889±5.966 | 0.000±0.000 | 0.000±0.000 | 5.845±5.757 |
| Medicine/Simple | 94.609±2.964 | 0.014±0.000 | 0.000±0.000 | 7.818±9.852 |

The WelshADF test revealed significant differences in AADPs among groups (92.45% variance explained), with search engines being the most important explanatory variable (WJ = 69265.22, df = 3, p < 0.001). Effects of the search topic (WJ = 8.49, df = 1, p = 0.005), keyword complexity (WJ = 71.71, df = 1, p < 0.001), the interaction of search topic and keyword complexity (WJ = 20.40, df = 1, p < 0.001), and their combination with search engine (Search engine × Topic: WJ = 11959.03, df = 3, p < 0.001, Search engine × Keyword complexity: WJ = 61790.69, df = 3, p < 0.001) on the outcome were all significant. The effect of browsers used was not significant, either alone (WJ = 0.06, df = 2, p = 0.941) or as a covariant of search engine choice (WJ = 0.29, df = 6, p = 0.943). Cache, whether it was emptied or not, did not have a significant effect, either in its own or as a covariant (Fig 1, Supplementary Information 1, Supplementary Information 2-3). In spite of not being a significant predictor in the entire dataset, both browser and cache showed a tendency to influence the outcome of the Google Scholar results. None of these influenced the search platforms with a background database. There were no differences in search results when using Web of Science, PubMed and Scopus but different machines at the same location but Google Scholar sometimes produced different results.

**Fig 1.**
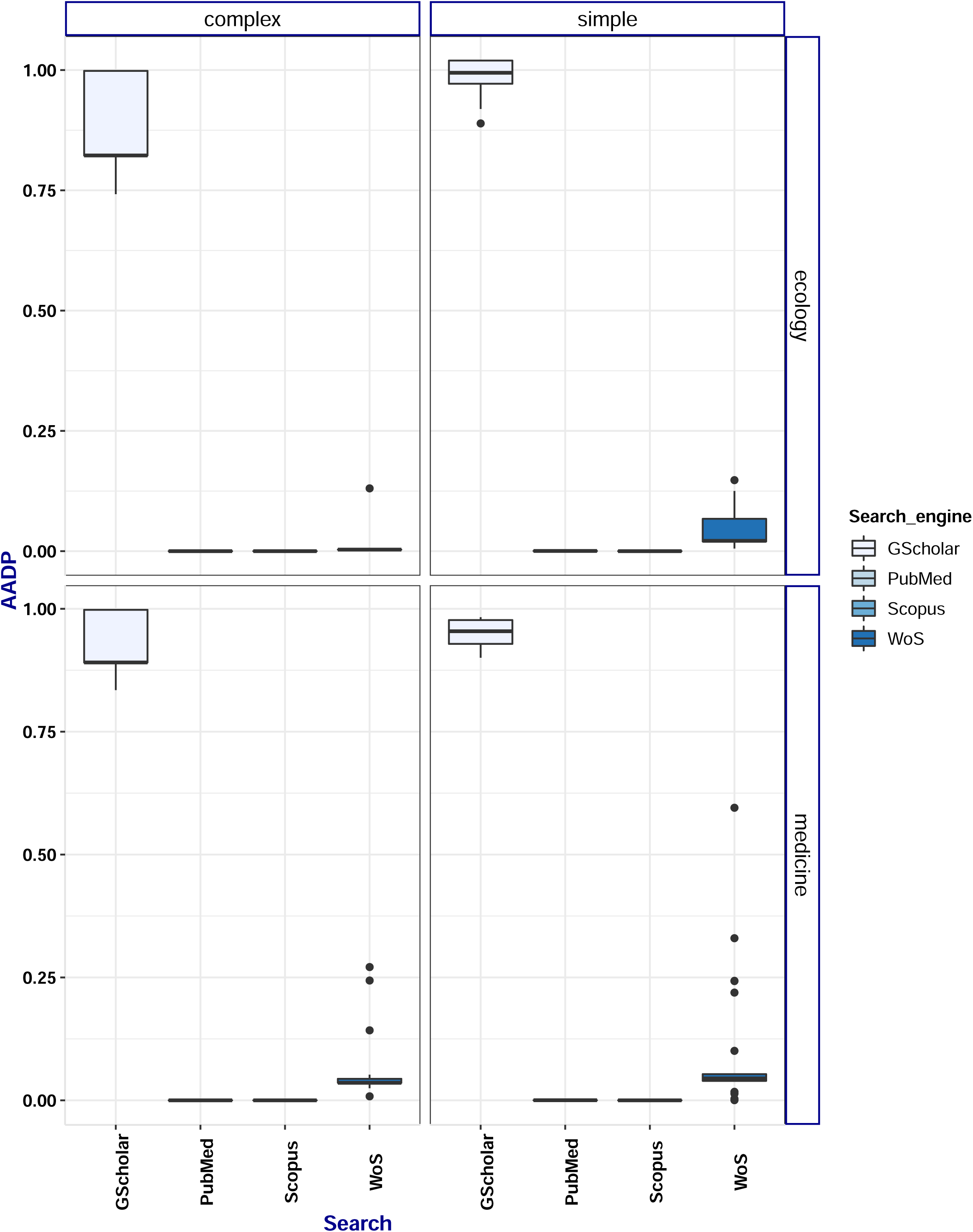
Average absolute deviation proportions (AADP) of hit numbers.

AADPs are grouped by searched platforms, and separated by keyword complexity (complex, simple), and research area (ecology, medicine).Boxes represent interquartile range (IQR), with median AADP values represented as a thick horizontal band. Whiskers extend from Q1-1.5IQR to Q3+1.5IQ. Abbreviated search platforms: GScholar – Google Scholar, WoS – Web of Science.

The multivariate analysis run on the first twenty papers collected from each search revealed significant differences among the search engines (p = 0.01) but did not show a significant influence on browser choice or cache state. Areas of convex hulls defined by these ‘paper-communities’ (see Methods) of the first twenty hits were zero for Scopus only, and they were the largest for Google Scholar (Table 3). When PubMed and Web of Science datasets were split by their update time, hulls for both PubMed subsets became zero but remained greater than zero for Web of Science. Distance measures showed an analogous pattern; they were zero for Scopus, indicating no difference between the first twenty papers, and deviated from zero for all other platforms (Fig 2). After correcting for the database update, only Web of Science and Google Scholar hulls remained significantly greater than zero.

**Table 3.** Areas of complex hulls for each search engines, separated by terms of topic and complexity.

| Topic/Complexity | GScholar | PubMed | Scopus | WoS |
| --- | --- | --- | --- | --- |
| Ecology/Complex | 491.90 | 0.00 | 0.00 | 0.00 |
| Ecology/Simple | 322.24 | 490.37 | 0.00 | 8.82 |
| Medicine/Complex | 476.45 | 4.99 | 0.00 | 0.02 |
| Medicine/Simple | 625.03 | 428.56 | 0.03 | 41.81 |

**Fig 2.**
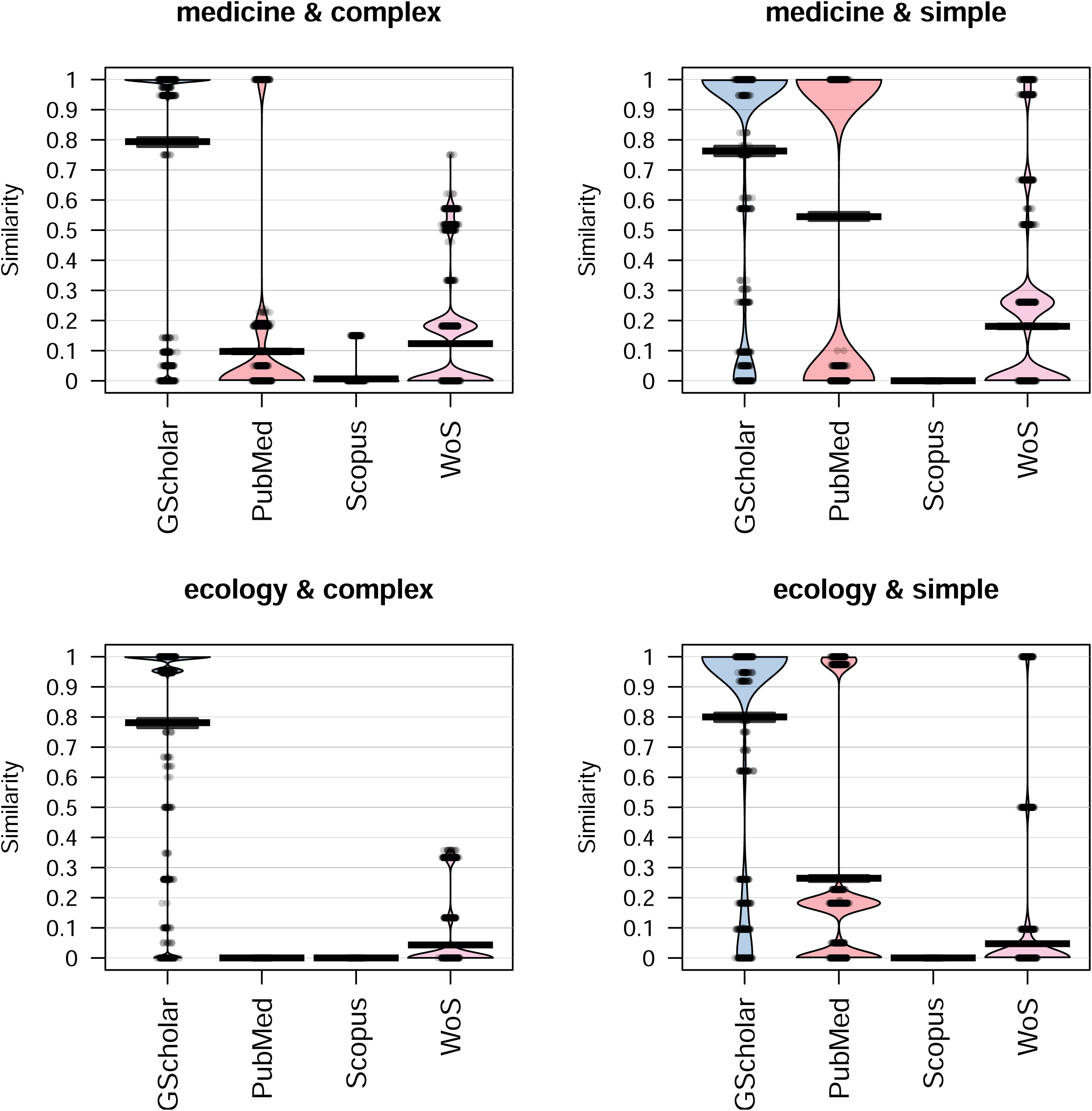
Average similarities of the first twenty papers within each search engine-topic-keyword complexity group, for each search platform.

Similarities were calculated based on binary matrices, using Jaccard distances. Median similarities are indicated with a thick black line on the pirate plots. Abbreviated search platforms: GScholar – Google Scholar, WoS – Web of Science.

## Discussion

In this study, we identified a shortcoming of scientific search platforms that can decrease the transparency and repeatability of the synthesis of quantitative evidence synthesis relying on database searches. Hence, the creditability and reliability of the conclusions drawn from these syntheses may be compromised.

Our results showed significant differences in search platform consistency in terms of both the number of hits (the size of the body of available evidence) and its composition when identical search terms were queried at different geographic locations. We found that PubMed and Scopus had high consistencies, whilst Google Scholar and Web of Science were not consistent in the number of hits they returned. Google Scholar provided the greatest number of hits for every search, it also proved to be the least consistent among different search runs, varying greatly in the number of hits, i.e. the total number of papers. Contrarily, the composition of the evidence collected, characterized by the first twenty papers it returned, was relatively consistent. Web of Science, however on a lower magnitude, showed similarly poor consistency in terms of the number of hits returned from identical searches initiated from different locations. Both the hit numbers and the returned list of articles from Scopus searches were consistent. PubMed varied in hit numbers and had great dissimilarities among the returned sets of papers, especially in those related to more general searches that necessarily had more hits. These dissimilarities were likely due to a database update that happened during our search exercise. Indeed, data showed that 0, 6, 10, 25 papers (complex ecology, complex medicine, simple ecology, and simple medicine terms, respectively) were added to the database during the course of this worldwide exercise. Since the papers listed were ordered according to their time of inclusion in the dataset, the first 20 collected papers would greatly differ and especially the larger values in the newly added articles can cause a disproportionally large effect on the similarity of the 20 collected papers. Once the differences before and after database update were accounted for, PubMed showed no deviation either in the number of returned papers or the list of the first 20 listed papers. A similar change in the dataset happened with Web of Science during our search, but differences remained even after correcting for the update. This suggests that discrepancies were caused by other sources, such as geographic locations. Overall, in our tests, Scopus and PubMed proved to be the most consistent databases, and Web of Science and Google Scholar produced highly inconsistent results.

Although we could not thoroughly decipher the influence of browser or cache on the search results, there was an indication that these factors only affected Google Scholar outcomes. Google Scholar is known to optimize search hits according to the search history of its users, thus, even the differences between browsers are likely to be the results of participants’ previous browser use, and therefore different cache contents in different browsers.

While the disadvantages of the inconsistencies in Google Scholar search results have been repeatedly illustrated[18,19], the similar behavior from Web of Science has only recently been reported[13] but in neither case was the variability estimated nor were the potential solutions discussed. Given the widespread use of Web of Science, neglecting this discrepancy can mislead scientists when drawing conclusions from their evidence synthesis, when the body of evidence was collected by Web of Science searches alone. The use of only one database is generally discouraged[5], and although some authors mainly target Google Scholar-based reviews[18,20], it is clear here that relying on Web of Science alone, or another single source, may lead to missing data or can make data-synthesis studies irreproducible. In spite of the recommendations of the need to use multiple sources for such studies (see the PRISMA statement[4]), a rapid scan of 20 recent papers in leading journals showed that recent, potentially highly cited, ecology-related systematic reviews still used Web of Science as their only search engine (Supplementary Information 4). In the light of the fact that using inadequate databases/search engines makes systematic reviews unreliable, our findings are concerning.

There are various means of overcoming this issue:

a. Researchers conducting systematic reviews should be aware of this potential problem, and be explicit about the methodology they use to ensure sufficient consistency and replicability. A detailed description should be included on the search engines used (ideally more than one), search dates, the exact search strings, as well as whether the same search was replicated by more than one person. As our study showed, the location from which the search was conducted should also be reported, preferably along with the IP address of the computer and the locality/institution the queries were initiated from. The exact time of the search or the time window of the query are also essential. The holdings of databases, however, are not constant, historical records can be added over time, and, therefore, queries even within a clearly limited time period can deliver different result sets. Thus, reporting the time window of the queries can provide only a partial solution.
b. The use of adequate search engines for a particular task should be an important consideration. All of the large databases have different strengths; Google Scholar searches grey literature, Web of Science has the largest (combined) dataset and, as our study confirmed, that Scopus and PubMed are the most consistent. Moreover, some databases may be more suitable for collecting information on a particular topic or have a greater historical coverage than others[14].
c. Providers of scientific search platforms should consider opening their search code and moving their paywalls to make reference lists publicly available[21], thus contributing to search consistency, and hence, scientific repeatability. Particularly Web of Science, as the most commonly used search engine, should act on making its search hits equally reachable to all users and, rather than *a priori* filtering them according to the institutions’ paywall, restrict access only *after* the primary result set has been provided to the user.
d. Google Scholar, on the other hand, should open its computer code to allow researchers to understand how hit lists are generated and how results are ordered. Google Scholar has been criticized by the scientific community for the obscurity of its search algorithms[22]. Although we acknowledge that this can be against business policies for some companies, we argue that compromises must be made for the sake of research integrity and scientific rigor.
e. Providing well-documented, standard application programming interfaces (APIs) and generating unique identifiers for searches, combining search term, result list, search time and location, and additional metadata (e.g. computing environment) is required. Using an API for standardized searches would be particularly beneficial for searches using Google Scholar that shows a strong dependence on the computing environment. Although this solution could control for a great deal of variation derived mostly from computing background and would be able to keep detailed records on the metadata of the searches, it also brings up novel challenges. Firstly, APIs can be more complex to use than simple web interfaces that may discourage users to use them. Moreover, collecting detailed data about search locations, or even computing environment, raises both security and privacy concerns. Finally, storing individual searches along with the necessary metadata may be resource heavy over a long period of time, which is likely to increase maintenance costs, and therefore the subscription fees, of these services.

Should these steps towards ensuring repeatability not happen, the critical voices to web-based systematic reviews can claim unreliability of this method[11]. Given that the systematic review methodology was originally developed to handle contentious issues with various, often conflicting bodies of evidence[5], this is a critical issue. This matter can only be exacerbated by the appearance of automatic systematic reviews, relying on artificial intelligence[23].

Despite the limited number of institutions that participated in this exercise, and the overrepresentation of Europe, the lack of contribution from African, South American and other Asian countries, we found, even within the European countries, variation among the numbers of search hits. This suggests that adding more countries would have led to even greater variability in the resulting datasets. It may be interesting to test a wider range of search platforms and subjects to gain further understanding of the level of reliability of various systems and collect reliable knowledge on their strengths and weaknesses.

Since, the original set of raw data input can significantly alter/skew the output of the study and, in the age of big data, studies on already published results are becoming more common, an unbiased and timely way of data extraction is needed. At present, updating systematic reviews using precisely repeated methodology is impossible[24]; hence a clear decision map on the advantages and disadvantages of particular databases and search engines should be drawn to ensure the integrity of publication-based studies.

## Materials and methods

### Queried databases

Three major scientific databases, PubMed, Scopus, and Web of Science, and Google Scholar, as the most used and largest scientific were used in this study. Although Google Scholar is markedly different from the other three traditionally used databases, both in business politics and search method[14,18], the increasing use of this search engine [20] justifies its inclusion in the study. The main differences between these databases have been catalogued and reviewed by Falagas et al.[14].

#### PubMed

(https://www.ncbi.nlm.nih.gov/pubmed) is a freely available scientific database, focusing mostly on biomedical literature, which holds ca. 28 million citations covering a variety of aspects of life sciences (https://www.ncbi.nlm.nih.gov/books/NBK3827/#pubmedhelp. PubMed_Coverage, accessed 15/08/2018). It was developed and is being maintained by the National Center for Biotechnology Information.

#### Scopus

currently owned by the Elsevier group, contains bibliographic data of over 1.4 billion publications dating back to 1970. It indexes ca. 70 million items and 22,500 journals from 5,000 publishers (https://www.elsevier.com/solutions/scopus/how-scopus-works/content, accessed: 17. August 2018).

#### Web of Science

(https://webofknowledge.com) is the oldest scientific database, owned by the Clarivate Analytics (previously Thomson Reuters). Web of Science, running under its current name since 1997, is the successor of the first scientific citation database, the Science Citation Index, which was launched in 1964. It currently indexes 34,200 journals, books and proceedings, and, as of the last update, on 26 August 2018, it covers 151 million records altogether and over 71 million in its Core Collection (https://clarivate.libguides.com/webofscienceplatform/coverage). Currently it also includes Zoological Records, CABI Abstracts, and a number of other, formerly independent databases.

#### Google Scholar

(https://scholar.google.com) is a free online tool, the sub-site of the search mogul Google Inc., which is particularly designed for scholarly searches. Whilst Google Scholar has been often criticized for not sharing its search algorithms, for its untraceable way of ordering search hits, and for the inclusion of material from non-scholarly sources in its research hits[18,19,25], it has been playing an increasing role in daily lives of scientists since its launch in 2004[20,26]. It is also estimated to include 160 million individual scientific publications in 2014[27] and to be the fastest growing resource for scientific literature[28]. Its usefulness, however, for systematic reviews and meta-analyses has been debated[16,18,19]

### Web searches

In order to investigate the reproducibility of scientific searches in the four major search platforms, we generated keyword expressions (search strings) with two complexity levels using keywords that focused on either an ecological or a medical topic and ran standardized searches from various locations in the world (see below), all within a limited timeframe.

Simple search strings contained only one main keyword, whereas complex ones contained both inclusion and exclusion criteria for additional, related, keywords and key phrases (i.e. two-word expressions within quotation marks). Wildcards (e.g. asterisks) and logical operators were used in complex search strings. The main common keyword for ecology was “ecosystem” and “diabetes” was used for the medical topic. Search language was set to English in every case, and only titles, abstracts and keywords were searched.

Since different search engines use slightly different expressions for the same query, exact search terms were generated for each search (Table 4).

**Table 4.** Search strings for each keyword complexity and topic, adjusted according to the search engines.

| Search engine | <u>Ecology</u> | Simple search string | <u>Medicine</u> | Simple search string |
| --- | --- | --- | --- | --- |
|  | Complex search string |  | Complex search string |  |
| **GScholar** | "ecosystem service"+"promoting"+"crop"-"livestock" | "ecosystem services" | "diabetes"+"sugar"+"fructose"-"saccharose" | "diabetes mellitus" |
| **PubMed** | "ecosystem service"[Title/Abstract] AND "promoting" AND "crop"[Title/Abstract] NOT "livestock"[Title/Abstract] AND "english"[Language] | "ecosystem services"[Title/Abstract] AND "english"[Language] | "diabetes"[Title/Abstract] AND "sugar" AND "fructose"[Title/Abstract] NOT "saccharose"[Title/Abstract] AND "english"[Language] | "diabetes mellitus"[Title/Abstract] AND "english"[Language] |
| **Scopus** | TITLE-ABS-KEY ("ecosystem service" AND "promoting" AND "crop" AND NOT "livestock") AND (LIMIT-TO (LANGUAGE, "English")) | TITLE-ABS-KEY ("ecosystem services") AND (LIMIT-TO (LANGUAGE, "English")) | TITLE-ABS-KEY ("diabetes" AND "sugar" AND "fructose" AND NOT "saccharose") AND (LIMIT-TO (LANGUAGE, "English")) | TITLE-ABS-KEY ("diabetes mellitus") AND (LIMIT-TO (LANGUAGE, "English")) |
| **WoS** | TS=("ecosystem service" AND "promoting" AND "crop" NOT "livestock") | TS=("ecosystem services") | TS=("diabetes" AND "sugar" AND "fructose" NOT "saccharose") | TS=("diabetes mellitus") |

Searches were conducted on one or two machines at 12 institutions in Australia, Canada, China, Denmark, Germany, Hungary, UK, and the USA (Supplementary Information 5), using the three main browsers (Mozilla Firefox, Internet Explorer, and Google Chrome). Searches were run manually (i.e. no APIs were used) according to strict protocols, which allowed to standardize search date, exact search term for every run, and data recording procedure. Not all databases could have queried from every location: Google was not available in China, and Scopus was not available at some institutions (Supplementary Information 5). The original version of the protocol is provided in Supplementary Information 6. The first run was conducted at 11:00 Australian Eastern Standard Time (01:00 GMT) on 13 April 2018 and the last search run at 18:16 on 13 April 2018 Eastern Daylight Time (22:16 GMT). After each search the number of resulted hits was recorded and the bibliographic data of the first 20 articles were extracted and saved in a file format that the website offered (.csv, .txt). Once all search combinations were run and browsers’ cache had been emptied, the process was repeated. At four locations (Flakkebjerg, Denmark; Fuzhou, China; St. Catharines, Canada; Orange, Australia) the searches were also repeated on two different computers.

Results were collected from each contributor, bibliographic information was stripped out from the saved files, and was stored in a standardized database, allowing unique publications to be distinguished. If unique identifiers for individual articles were missing, authors, titles, or the combination of these were searched for, and uniqueness was double checked across the entire dataset.

For the rapid scan, if authors used Web of Science as the main search platform, and if search locations were reported, we chose the first twenty papers from a Google Scholar search (7 November, 2018) with the search term “systematic review” and “ecology”. Sites were restricted to sciencemag.org, nature.com, and wiley.com.

### Statistical analysis

To investigate how consistent the number of resulting hits from each search string (i.e. the combination of the search topic and keyword expression complexity) was for each of the search engines, *average absolute deviation* (AAD, i.e. the absolute value of the difference of the actual value and the mean) was calculated and expressed as a percentage of the mean of each group (‘*average absolute deviation proportion*’, AADP, i.e. search topic, search term complexity, and search engine). AADP was calculated using the equation:

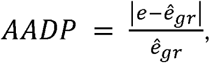

where *e* was the number of hits from one particular search and *ê_gr_* was the mean number of hits of pooled numbers from one topic and search term complexity combination and one search engine (e.g. complex ecological search expression queried using Scopus). This grouping was necessary because the number of hits substantially differed depending on these three factors. Since the aim of the study was not to compare the efficiency of different search engines, this grouping did not interfere with our analysis.

Normality of the data and homoscedasticity were tested using Kolmogorov-Smirnoff test and the Breusch Pagan test, respectively. These tests confirmed that neither the distribution of AADPs followed normal distribution, nor were the variances of residuals within each group homogenous. Indeed, the high number of zeroes resulted in a zero-inflated, an unbalanced beta distribution, as suggested by the *descdist*() function in the *fitdistrplus* R package[29], under an R programming environment[30].

AADP is expected to be zero in cases when search engines consistently give the same number of hits within groups, regardless where the search is initiated from, browser used, or whether the cache was emptied or not. Therefore, one-sided Wilcoxon signed rank tests were performed for the AADP values for each search engines within each group to test if they were significantly different from zero.

To address non-normality, unequal variances and control Type I error, non-parametric, Welch-James’s statistic with Approximate Degrees of Freedom (Welch ADF) was used to investigate the differences between search engine consistencies and to select the most influential factors driving these differences. This robust estimator uses trimmed means and winsorized variances to avoid biases derived from heteroscedasticity. Bootstrapping was used to calculate empirical p-values both for between group and pairwise comparisons[31], with the help of WelchADF R package[32].

Moreover, average similarities of the first twenty papers within each of the search engine-topic-keyword complexity groups were calculated based on binary matrices, in which rows corresponded to search runs from various institutions and computers, whilst columns contained individual papers. Due to its suitability for using binary data, Jaccard distance measures were applied for similarity calculations. Distance-based redundancy analysis (dbRDA, capscale() function) was used with the same similarity matrices to ordinate the resultant article collections in each search topic-keyword complexity group. Convex hulls of the points resulted from this ordination were then delimited for each search engine and their areas were calculated. Since similarities between article collections resulted from searches with a search engine giving consistently the same hits, regardless of search location, browser used, and cache content, should always be zero, the ideal size of these hulls would be also zero.

## Data availability statement

All data and computer code are deposited on the Open Science Framework (OSF) website and will be openly available for the readers through a stable URL or DOI upon acceptance.

## Acknowledgements

We are grateful for Dr. Mei Ling Huang (Brock University, St. Catharines, Canada) for her insightful comments on the statistical analysis. This work is supported by a grant of “111 project” in China. Gabor Pozsgai is supported by a postdoctoral fellowship by the State Key Laboratory of Ecological Pest Control for Fujian and Taiwan Crops. Arnold Móra was supported by the grants #20765-3/2018/FEKUTSTRAT and #TUDFO/47138/2019-ITM.

## Author contributions

Gábor Pozsgai and Geoff Gurr conceived the project. Gábor Pozsgai designed the experiment, and did the statistical analysis. Gábor Lövei, Gábor Pozsgai, Jie Zhang, and Wenwu Zhou performed the preliminary searches. All contributors were involved in running the searches and providing raw data in the given format. The first drafted version of the manuscript was prepared by Gábor Pozsgai. This draft was first edited by Gábor Lövei, Liette Vasseur, Geoff Gurr, Olivia Reynolds, and Minsheng You. All authors were included in editing the subsequent versions of the manuscript. Minsheng You funded the work.

## Competing interests

The authors declare no competing interest.

**Supporting Information** 1. The results of the Welch-James’s statistic with Approximate Degrees of Freedom. Significant (p < 0.05) relationships are highlighted with bold font.

**Supporting Information 2**. Average absolute deviation proportions (AADP) of hit numbers, grouped by searched platforms, and separated by grouped keyword complexity (complex, simple) – research area (ecology, medicine) and cache state. Boxes represent interquartile range (IQR), with median AADP values represented as a thick horizontal band. Whiskers extend from Q1-1.5IQR to Q3+1.5IQ. Abbreviated search platforms: GScholar – Google Scholar, WoS – Web of Science.

**Supporting Information 3**. Average absolute deviation proportions (AADP) of hit numbers, grouped by searched platforms, and separated by grouped keyword complexity (complex, simple) – research area (ecology, medicine) and browser type. Boxes represent interquartile range (IQR), with median AADP values represented as a thick horizontal band. Whiskers extend from Q1-1.5IQR to Q3+1.5IQ. Abbreviated search platforms and browsers: GScholar – Google Scholar, WoS – Web of Science, Chrome – Google Chrome, IE – Internet Explorer, Mozilla – Mozilla Firefox.

**Supporting Information 4**. The list of papers used in the rapid screen and the results showing how many different search platforms were used and whether or not the date, search location and browser were indicated.

**Supporting Information 5**. Names and affiliations of contributors and list of scientific search platforms accessed during the search exercise.

**Supporting Information 6**. The exact protocol which was circulated to contributors, describing how searches should be performed and how data should be saved.

## References

1. McMullin E. The Impact of Newton’s Principia on the Philosophy of Science. Philos Sci. 2001;68: 279–310. doi: 10.1086/392883

2. Baker M. 1,500 scientists lift the lid on reproducibility. Nature. 2016;533: 452–454. doi: 10.1038/533452a

3. Cook DJ, Mulrow CD, Haynes RB. Systematic reviews: Synthesis of best evidence for clinical decisions. Ann Intern Med. 1997;126: 376–380. doi: 10.7326/0003-4819-126-5-199703010-00006

4. Moher D, Liberati A, Tetzlaff J, Altman DG. Academia and Clinic Annals of Internal Medicine Preferred Reporting Items for Systematic Reviews and Meta-Analyses: The PRISMA Statement. Annu Intern Med. 2009;151: 264–269. doi: 10.1371/journal.pmed1000097

5. Higgins J, Green S. Cochrane Handbook for Systematic Reviews of Interventions. Chichester, UK: The Cochrane Collaboration; 2008. Available: http://www.cochrane-handbook.org/

6. Bornmann L, Mutz R. Growth rates of modern science: A bibliometric analysis based on the number of publications and cited references. J Assoc Inf Sci Technol. 2015;66: 2215–2222. doi: 10.1002/asi.23329

7. Pain E. How to keep up with the scientific literature. Science Careers. 30 Nov 2016. doi: 10.1126/science.caredit.a1600159

8. Landhuis E. Scientific literature: Information overload. Nature. 2016;535: 457–458. doi: 10.1038/nj7612-457a

9. Gurevitch J, Koricheva J, Nakagawa S, Stewart G. Meta-analysis and the science of research synthesis. Nature. 2018;555: 175–182. doi: 10.1038/nature25753

10. Clarke M, Horton R. Bringing it all together: Lancet-Cochrane collaborate on systematic reviews. Lancet. 2001;357: 1728. doi: 10.1016/S0140-6736(00)04934-5

11. Ioannidis JPA. The Mass Production of Redundant, Misleading, and Conflicted Systematic Reviews and Meta-analyses. Milbank Q. 2016;94: 485–514. doi: 10.1111/1468-0009.12210

12. Garg AX, Hackam D, Tonelli M. Systematic review and meta-analysis: When one study is just not enough. Clin J Am Soc Nephrol. 2008;3: 253–260. doi: 10.2215/CJN.01430307

13. Gusenbauer M, Haddaway NR. Which Academic Search Systems are Suitable for Systematic Reviews or Meta□Analyses? Evaluating Retrieval Qualities of Google Scholar, PubMed and 26 other Resources. Res Synth Methods. 2019; jrsm.1378. doi: 10.1002/jrsm.1378

14. Falagas ME, Pitsouni EI, Malietzis GA, Pappas G. Comparison of PubMed, Scopus, Web of Science, and Google Scholar: strengths and weaknesses. FASEB J. 2007;22: 338–342. doi: 10.1096/fj.07-9492LSF

15. Gavel Y, Iselid L. Web of Science and Scopus: a journal title overlap study. Online Inf Rev. 2008;32: 8–21. doi: 10.1108/14684520810865958

16. Boeker M, Vach W, Motschall E. Google Scholar as replacement for systematic literature searches: Good relative recall and precision are not enough. BMC Med Res Methodol. 2013;13. doi: 10.1186/1471-2288-13-131

17. Jaccard P. The distribution of the flora in the alpine zone. New Phytol. 1912;11: 37–50. doi: 10.1111/j.1469-8137.1912.tb05611.x

18. Jacsó P. Google Scholar revisited. Online Inf Rev. 2008;32: 102–114. doi: 10.1108/14684520810866010

19. Jacsó P. As we may search – Comparison of major features of the Web of Science, Scopus, and Google Scholar citation-based and citation-enhanced databases. Curr Sci. 2005;89: 1537–1547. Available: http://muse.jhu.edu/content/crossref/journals/library_trends/v056/56.4.jacso.html

20. Haddaway NR, Collins AM, Coughlin D, Kirk S. The role of google scholar in evidence reviews and its applicability to grey literature searching. PLoS One. 2015;10: 1–17. doi: 10.1371/journal.pone.0138237

21. Shotton D. Funders should mandate open citations. Nature. 2018. doi: 10.1038/d41586-018-00104-7

22. van Dijck J. Search engines and the production of academic knowledge. Int J Cult Stud. 2010;13: 574–592. doi: 10.1177/1367877910376582

23. Beller E, Clark J, Tsafnat G, Adams C, Diehl H, Lund H, et al. Making progress with the automation of systematic reviews: Principles of the International Collaboration for the Automation of Systematic Reviews (ICASR). Syst Rev. 2018;7: 1–7. doi: 10.1186/s13643-018-0740-7

24. Garner P, Hopewell S, Chandler J, MacLehose H, Schünemann HJ, Akl EA, et al. When and how to update systematic reviews: Consensus and checklist. BMJ. 2016;354: 1–10. doi: 10.1136/bmj.i3507

25. Noruzi A. Google Scholar: The New Generation of Citation Indexes. Libri. 2005;55: 170–180. doi: 10.1515/LIBR.2005.170

26. Halevi G, Moed H, Bar-Ilan J. Suitability of Google Scholar as a source of scientific information and as a source of data for scientific evaluation—Review of the Literature. J Informetr. 2017;11: 823–834. doi: 10.1016/j.joi.2017.06.005

27. Orduna-Malea E, Ayllón JM, Martín-Martín A, Delgado López-Cózar E. Methods for estimating the size of Google Scholar. Scientometrics. 2015;104: 931–949. doi: 10.1007/s11192-015-1614-6

28. Larsen PO, von Ins M. The rate of growth in scientific publication and the decline in coverage provided by science citation index. Scientometrics. 2010;84: 575–603. doi: 10.1007/s11192-010-0202-z

29. Delignette-Muller ML, Dutang C. fitdistrplus: An R Package for Fitting Distributions. J Stat Softw. 2015;64: 1–34. Available: http://www.jstatsoft.org/v64/i04/

30. R Core Team. R: A Language and Environment for Statistical Computing. Vienna, Austria; 2012. Available: http://www.r-project.org/

31. Keselman HJ, Algina J, Lix LM, Wilcox RR, Deering KN. A generally robust approach for testing hypotheses and setting confidence intervals for effect sizes. Psychol Methods. 2008;13: 110–129. doi: 10.1037/1082-989X.13.2.110

32. Villacorta PJ. welchADF: Welch-James Statistic for Robust Hypothesis Testing under Heterocedasticity and Non-Normality. 2018. Available: https://cran.r-project.org/package=welchADF

