## Supporting information 1 for "A comparative analysis reveals irreproducibility in searches of scientific literature"

Supplementary Table 1. The results of the Welch-James's statistic with Approximate Degrees of Freedom. Significant (p < 0.05) relationships are highlighted with bold font.

|  | WJ statistics | Numerator DF | Denominator DF | P-value |
| --- | --- | --- | --- | --- |
| **Search engine** | **69265.22** | **3** | **86.24** | **> 0.001** |
| **Topic** | **8.49** | **1** | **64.98** | **0.005** |
| **Keyword complexity** | **71.71** | **1** | **64.98** | **> 0.001** |
| Browser | 0.06 | 2 | 43.30 | 0.941 |
| Cache | 0.12 | 1 | 64.98 | 0.729 |
| **Search engine × Topic** | **11959.03** | **3** | **86.24** | **> 0.001** |
| **Search engine × Keyword complexity** | **61790.69** | **3** | **86.24** | **> 0.001** |
| **Topic × Keyword complexity** | **20.40** | **1** | **64.98** | **> 0.001** |
| Search engine × Browser | 0.29 | 6 | 72.76 | 0.943 |
| Topic × Browser | 0.11 | 2 | 43.30 | 0.893 |
| Keyword complexity × Browser | 0.01 | 2 | 43.30 | 0.988 |
| Search engine × Cache | 0.11 | 3 | 86.24 | 0.952 |
| Topic × Cache | 0.01 | 1 | 64.98 | 0.934 |
| Keyword complexity × Cache | 0.07 | 1 | 64.98 | 0.799 |
| Browser × Cache | 0.02 | 2 | 43.30 | 0.985 |
| **Search engine × Topic × Keyword complexity** | **11955.76** | **3** | **86.24** | **> 0.001** |
| Search engine × Topic × Browser | 0.48 | 6 | 72.76 | 0.819 |
| Search engine × Keyword complexity × Browser | 0.06 | 6 | 72.76 | 0.999 |
| Topic × Keyword complexity × Browser | 0.02 | 2 | 43.30 | 0.984 |
| Search engine × Topic × Cache | 0.58 | 3 | 86.24 | 0.629 |
| Search engine × Keyword complexity × Cache | 0.10 | 3 | 86.24 | 0.962 |
| Topic × Keyword complexity × Cache | 0.05 | 1 | 64.98 | 0.831 |
| Search engine × Browser × Cache | 0.23 | 6 | 72.76 | 0.967 |
| Topic × Browser × Cache | 0.27 | 2 | 43.30 | 0.763 |
| Keyword complexity × Browser × Cache | 0.003 | 2 | 43.30 | 0.997 |
| Search engine × Topic × Keyword complexity × Browser | 0.33 | 6 | 72.76 | 0.921 |
| Search engine × Topic × Keyword complexity × Cache | 0.59 | 3 | 86.24 | 0.624 |
| Search engine × Topic × Browser × Cache | 0.18 | 6 | 72.76 | 0.983 |
| Search engine × Keyword complexity × Browser × Cache | 0.05 | 6 | 72.76 | 1.000 |
| Topic × Keyword complexity × Browser × Cache | 0.07 | 2 | 43.30 | 0.937 |
| Search engine × Topic × Keyword complexity × Browser × Cache | 0.06 | 6 | 72.76 | 0.999 |
