## Supplementary figures and images for "A comparative analysis reveals irreproducibility in searches of scientific literature"

### Supporting information 2

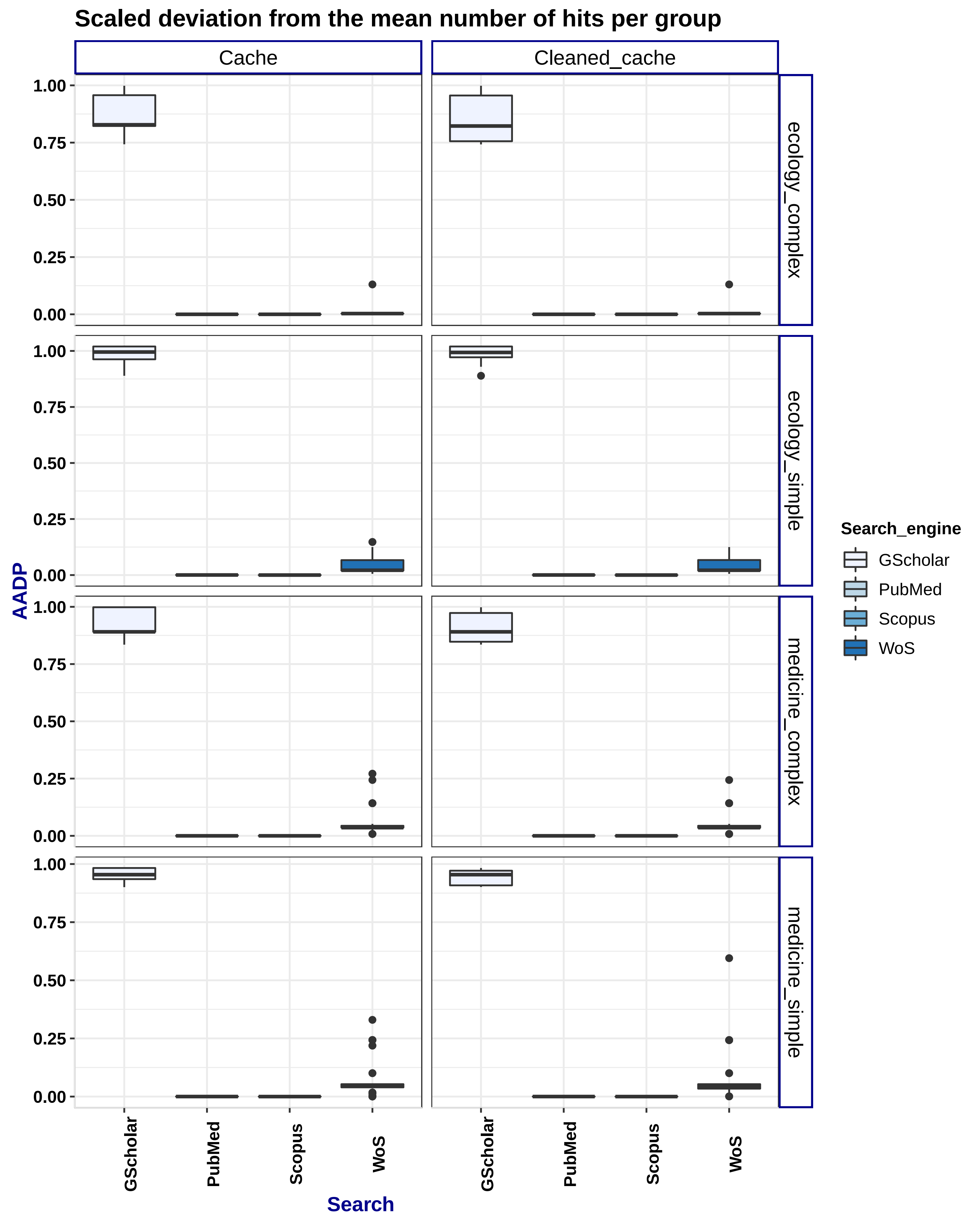

### Supporting information 3

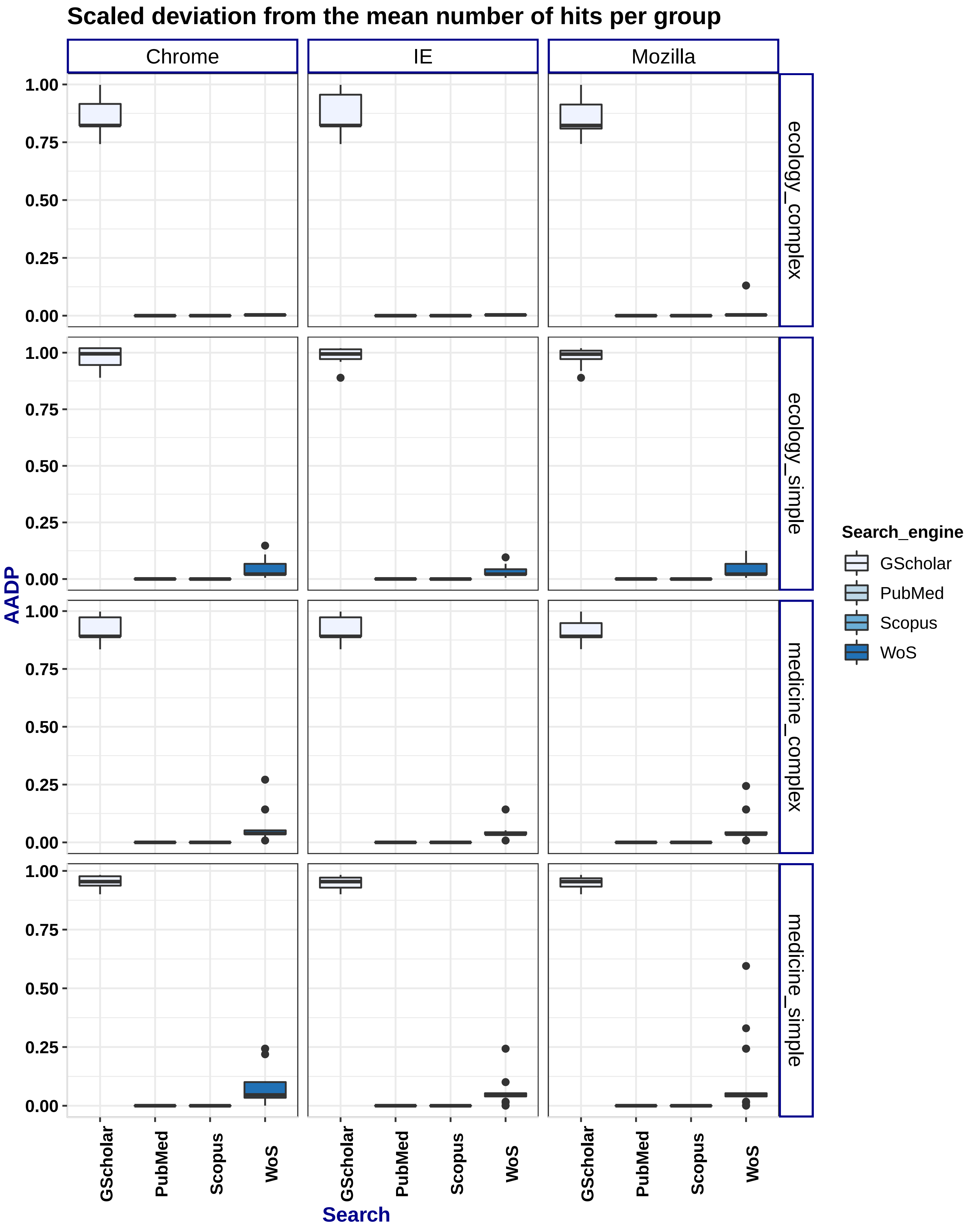
