## Supporting information 6 for "A comparative analysis reveals irreproducibility in searches of scientific literature"

Dear Colleagues,

Thanks for participating in our scientific search-exercise. In order to ensure consistency amongst searches you are asked to follow the protocol detailed below.

1. Make sure you have Google Chrome (GC), Mozilla Firefox (MF) and Internet Explorer (IE) installed on your computer
2. Make sure you have access to Web of Science (WoS), Scopus, PubMed and Google Scholar (GS). Even if you do not have access to all of these search engines you still can continue with the process but you have to mark in the data input sheet (see below) with an “NA” that that particular database was not available at your location.
3. Open GC, MF and IE.
4. Open four tabs in each with one of the search engines (WoS, Scopus, PubMed and GS).
5. Make sure that all the search engines are set as listed below
   a. Wos
      - Database is set to ‘Web of Science Core Collection’
      - You are on ‘Advanced search’
      - Language is set to ‘English’
      - ‘All years’ are searched
      - Hits are sorted according the date
   b. Scopus
      - Hits are sorted according the dates (‘Date(newest)’)
   c. PubMed
      - The database is set to PubMed (top left corner)
      - Hits are sorted according the dates (‘Most recent’)
   d. GS. Open the Setting by clicking on the three horizontal bars in the top left corner.
      - In the first tab (Search results), untick ‘Include patents’
      - Set results per page to 20 (Fig. 1)
      - Set your search language is on English only (Fig. 2)
      - On the main page untick ‘Include citations’ and ‘Include patents’
      - Sort your hits by date

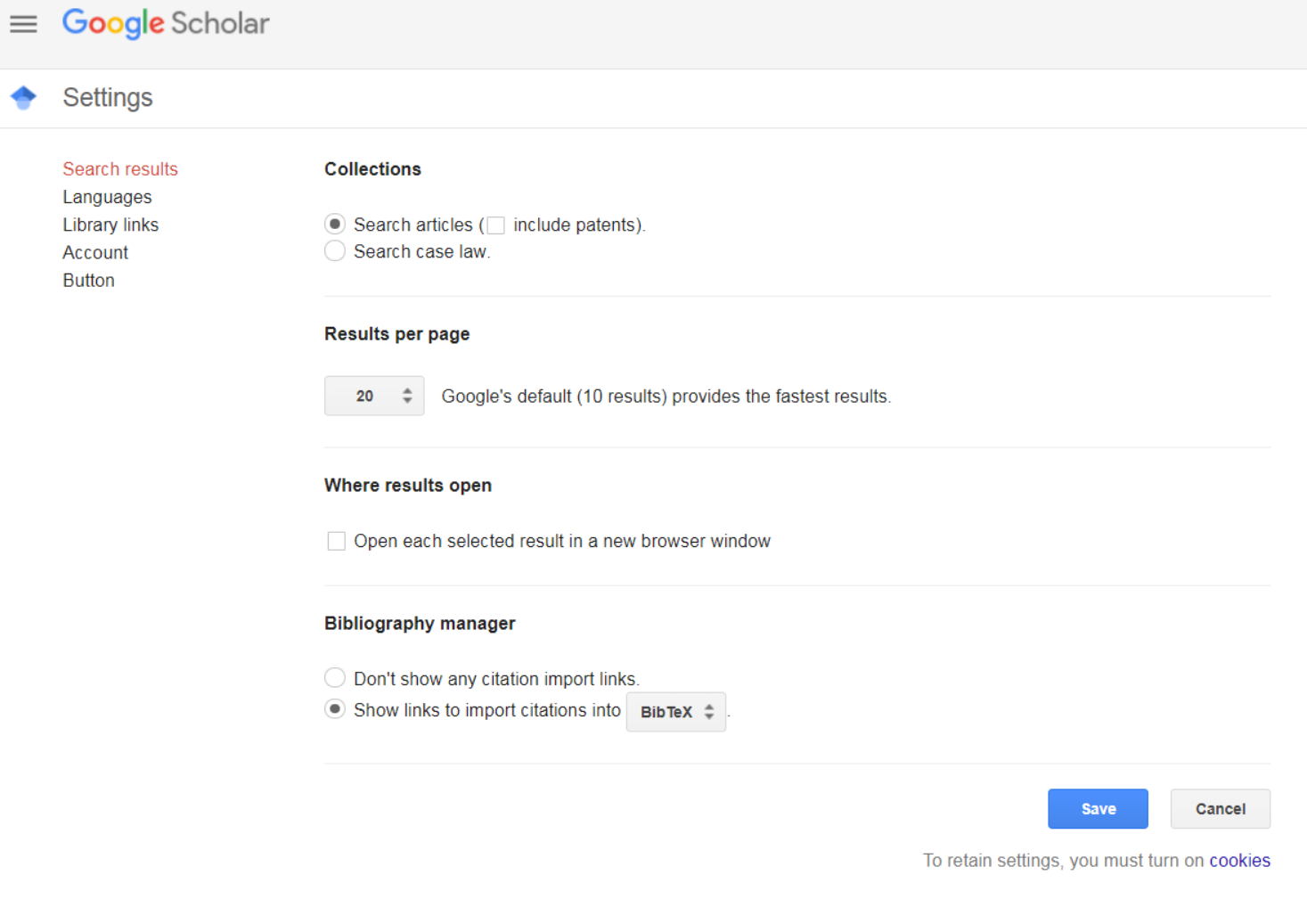

Figure 1.

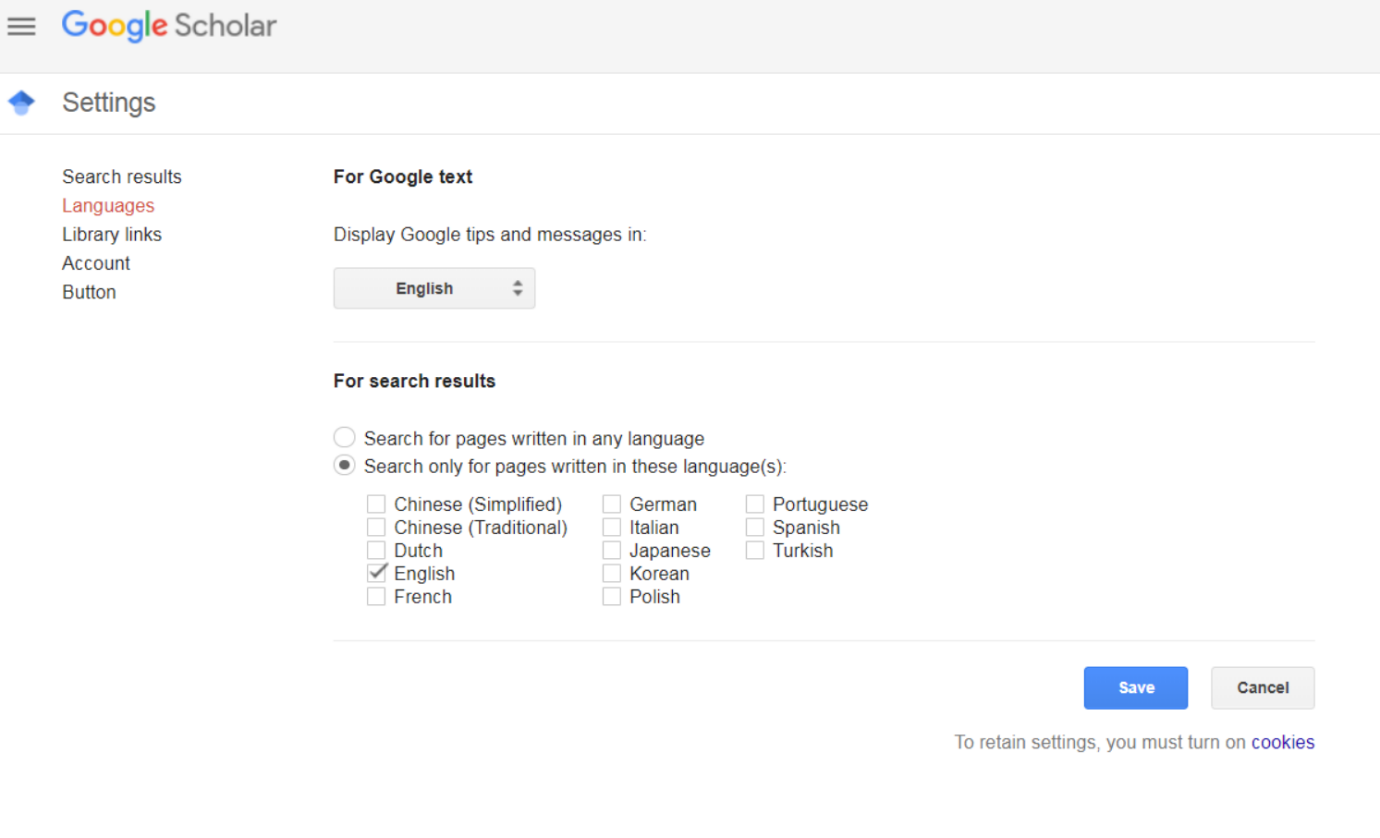

Figure 2.

6. Search process and saving data.
   a. Searches have to be run within the agreed timeframe.
   b. Please note the time when you start and stop running the searches.
   c. At each location searches with two keyword expression complexity, using four search engines in three browsers on two topics. Optimally, searches should be run on two separate machines, with and without the cache emptied (or cookies disabled) and repeated twice (different days, but the replication time should be agreed as well across contributors). This results in 96 searches for each machine, each replica (2-3 hours work).
   d. An excel file with all necessary setting and searchword combinations is circulated. Each row in the spreadsheet contains a keyword expression for each combination of expression complexity, search engines, browser, and keyword topic, and thus, each row translates to one search. Go through each row when you run the searches in order, by simply copying searchword expression from the cell and pasting it into the search field in your browser. Do not reorder this file!
   e. Start with searches that allow the use of cache (Unless you particularly disabled cache in your browser you do not need to do anything to allow this). Once these all have been completed, empty your cache in each browser (see below) and continue with searches that need cleaned cache.
   f. Number of hits should be recorded in the attached recording sheet for each search.
   g. **The first 20 hits should be saved** for each search with a file name comprising of the (abbreviated) name of the affiliating institute, indicating whether the search was run on the first or second computer, the number of replica and the number of row in the excel table to which the search is linked to. Output files will be saved in a .csv format (e.g. FAFU_2_1_97.csv = Fujian Agriculture and Forestry University, 2^nd^ computer, 1^st^ replica, running the last (97^th^) row in the excel table). File names, along with the number of hits resulted from the search should be recorded in the attached recording sheet. On some sites it is not possible to limit the number of saved outputs to 20. In these cases the nearest possible number, exceeding 20, should be chosen for the output. Results from different computers should be recorded in separates files.
7. Details for search engines
   a. WoS
      - Restart new session if it is necessary.
      - Copy and paste search term into the query field in advanced search.
      - Search history will show your most recent search on the top of the list, number of hits are indicated here. Record it in the excel file. You have to click on this number to bring up results (Fig. 3).
      - At the bottom of the page you can set the number of records you want to see. Set it 25 per page.
      - Make sure hits are sorted by date
      - Hit ‘Select page’ to mark every paper on the actual page
      - Roll down menu beside ‘Save to EndNote online’ and choose and choose ‘Save other file formats’
      - In the pop-up window mark records 1 to 20 and set the format to ‘Tab-delimited (Win, UTF-8)’ and hit ‘Send’ (Fig. 4)
      - Name the output file according to 6/f point, above and record name in the excel file.

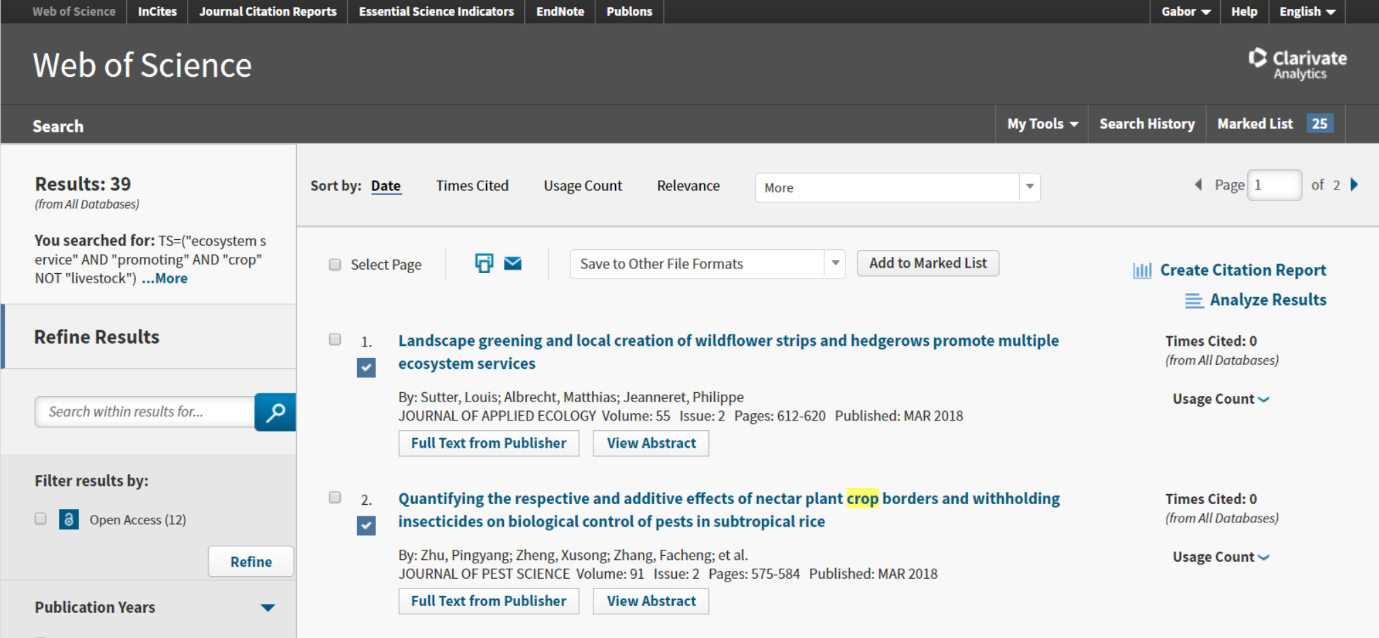

Figure 3.

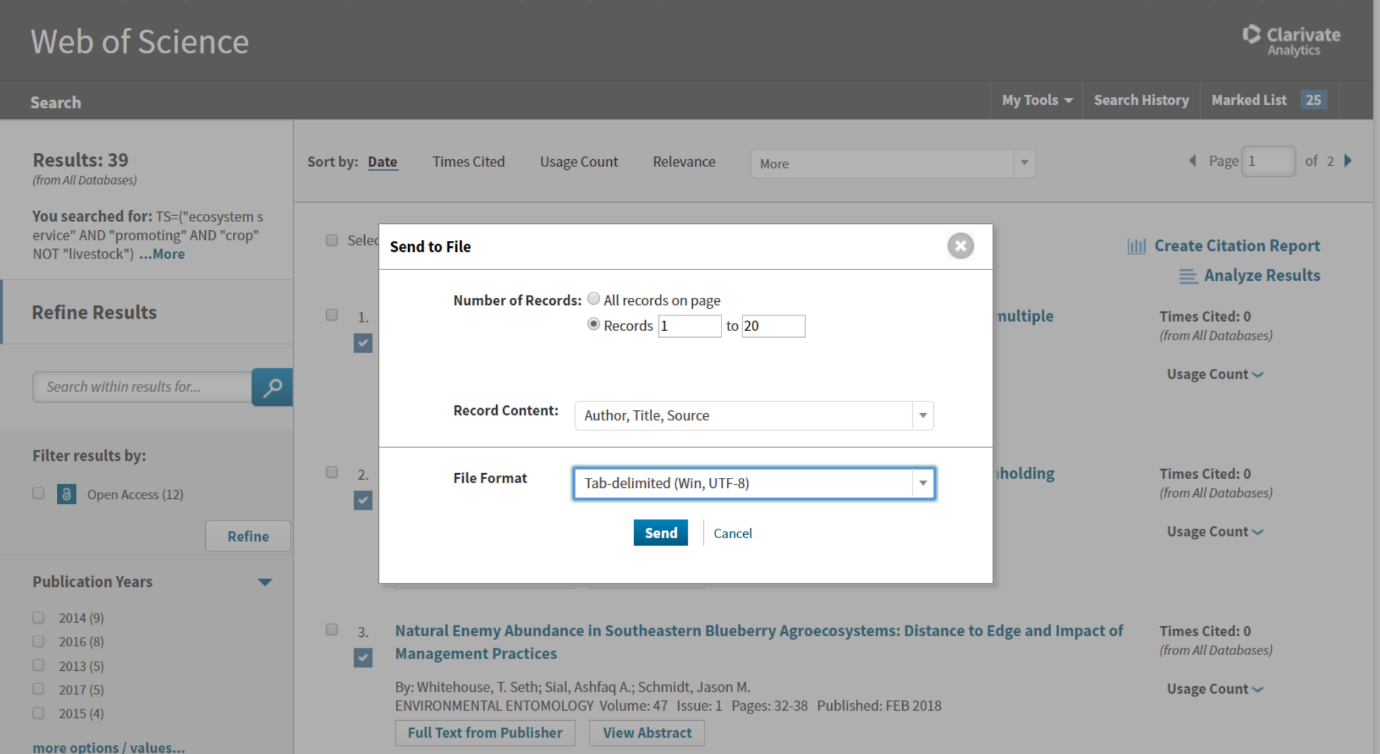

Figure 4.

b. Scopus
   - Copy and paste search term into the query field in advanced search.
   - The number of hits is indicated in the wide blue line on the top of the page. Please record this, in the excel file.
   - Make sure results are sorted according to date and 20 hits are shown on one page. Select all 20 hits and click on ‘Export’ (Fig. 5.).
   - Select ‘CSV’ in the pop-up window, and save citation information. Please also select ‘PubMed ID’ (Fig. 6.). Record file name in the excel file.

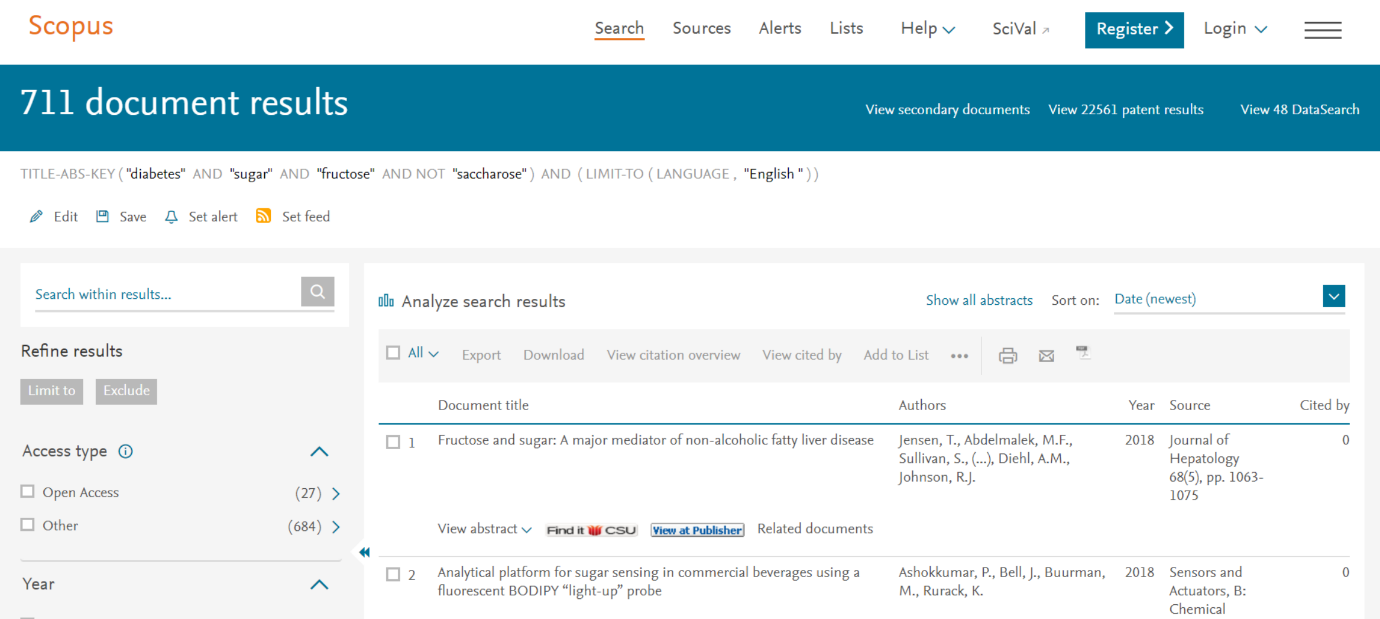

Figure 5.

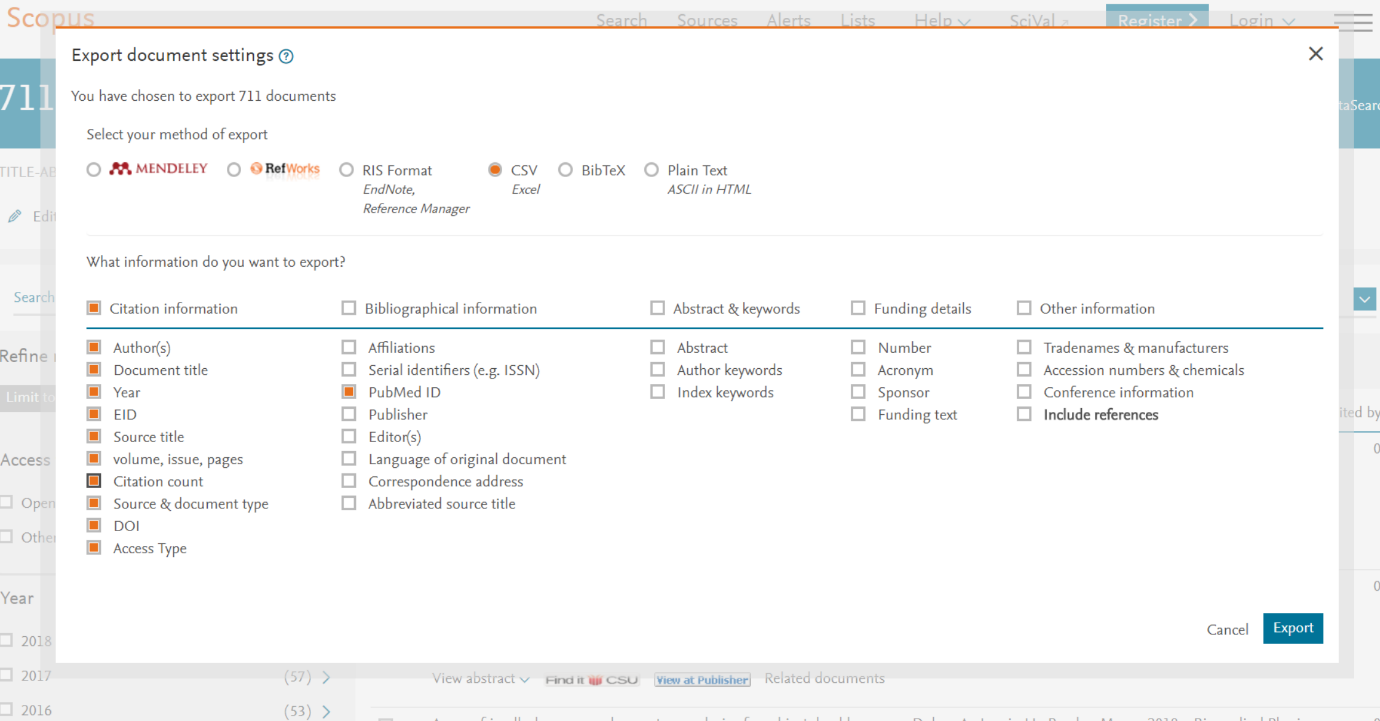

Figure 6.

c. PubMed
   - Copy and paste search term into the query field on the <https://www.ncbi.nlm.nih.gov/pubmed> site.
   - Make sure records are sorted by date, with the most recent ones first, and that the page shows 20 records.
   - Number of hits can be found on the right side of the page, under the ‘Recent activity’ title (Fig. 7). Please record this.
   - Select 20 hits (or all, if the number of hits is lower than 20) and click on ‘Send to’.
   - Select file, format CSV, and make sure the order is set to ‘Most recent’ (Fig. 8). Record the file name in the excel file.

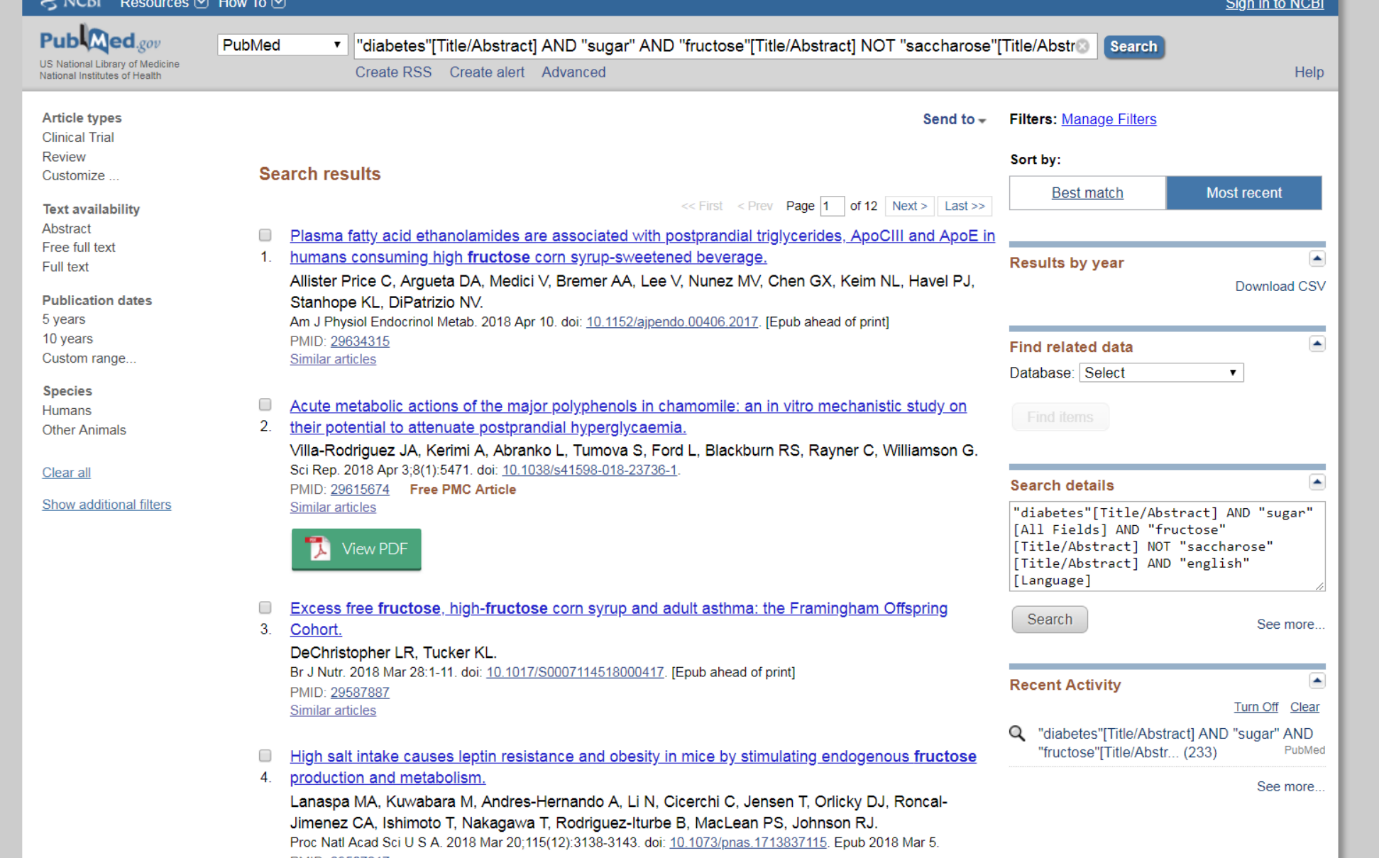

Figure 7.

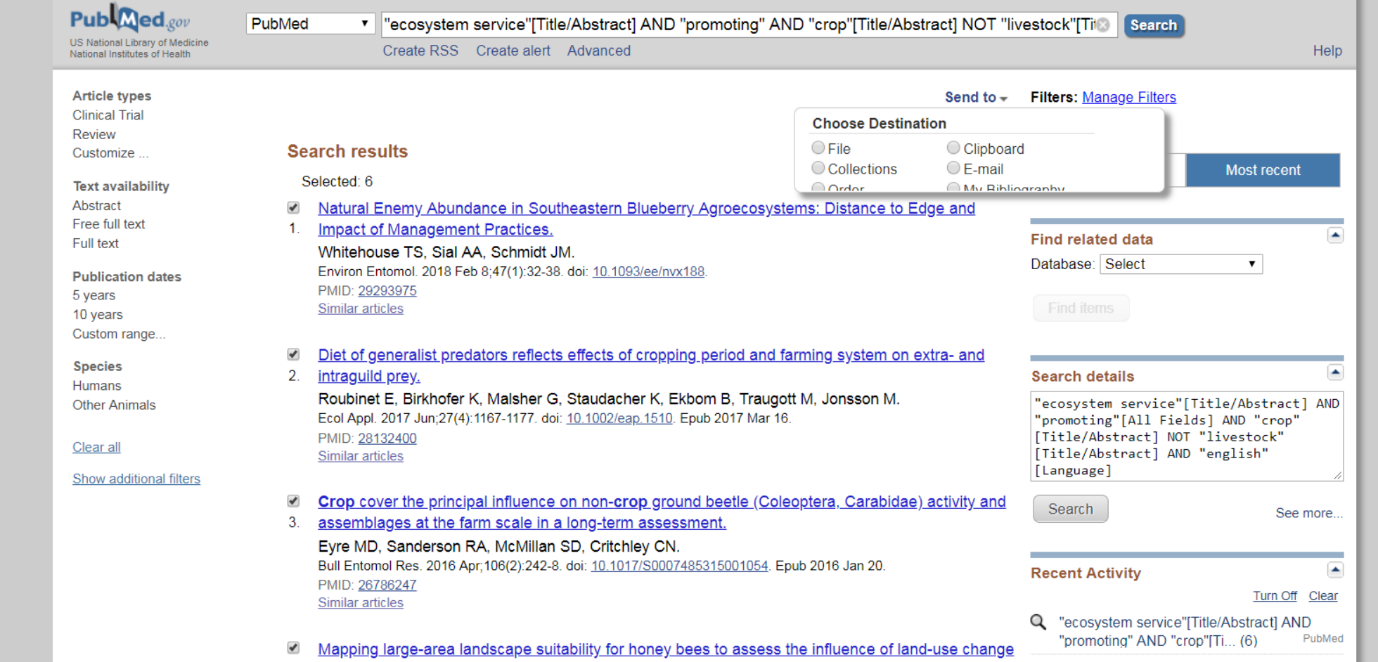

Figure 8.

d. Google Scholar
   - Sign into google
   - Make sure Google Scholar is set as it is indicated in 5/d
   - Copy and paste search term into the query field onto search box
   - Number of results are indicated below the search box
   - Make sure hits are ordered by date and mark the first 20 hits by clicking on the stars below the titles. These will be added to your google library now (Fig. 9).
   - Go to the ‘My library’, tick those 20 records you have just added and click on the ‘Export’ icon. Select CSV and save. Record file number in the Excel file (Fig. 10).

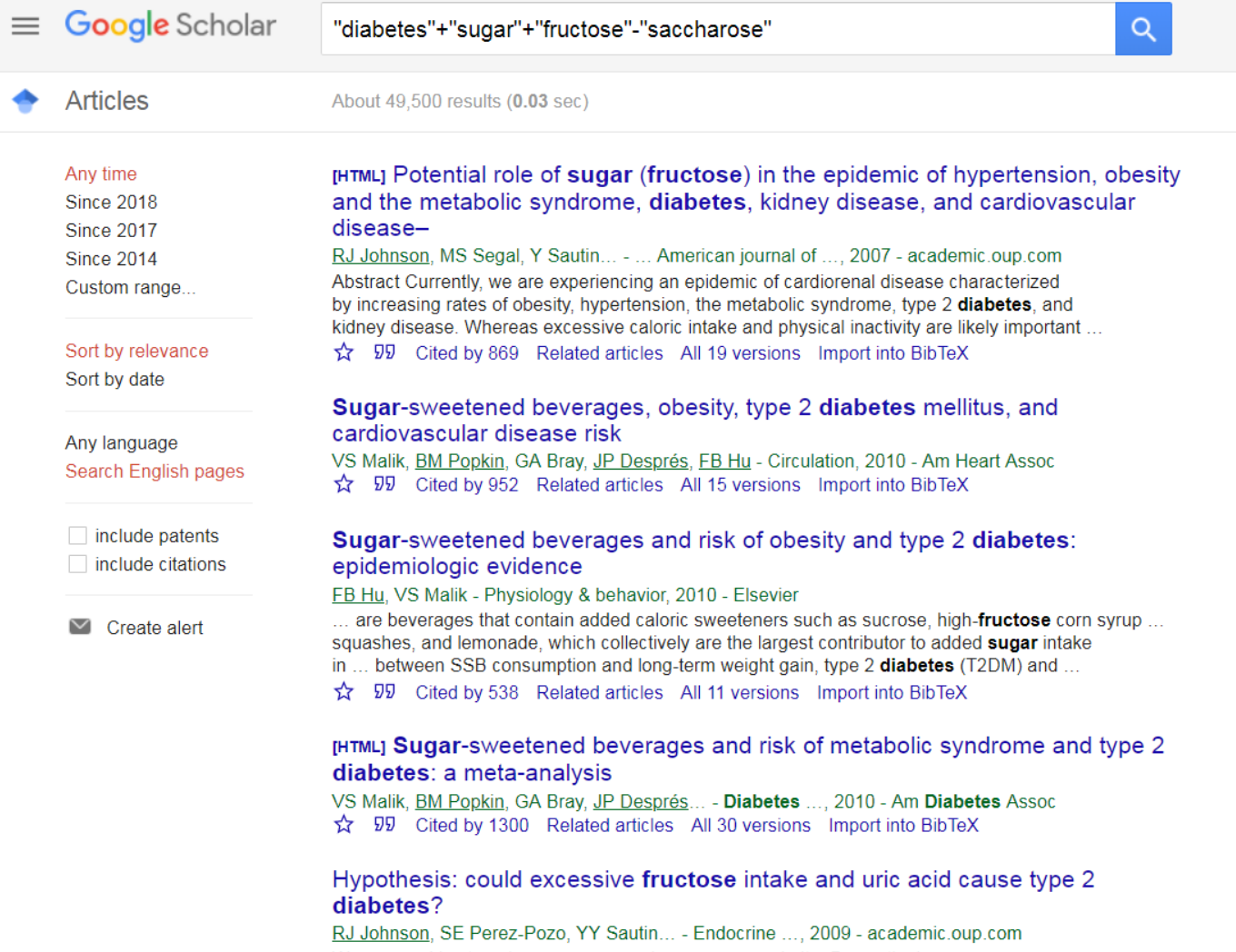

Figure 9.

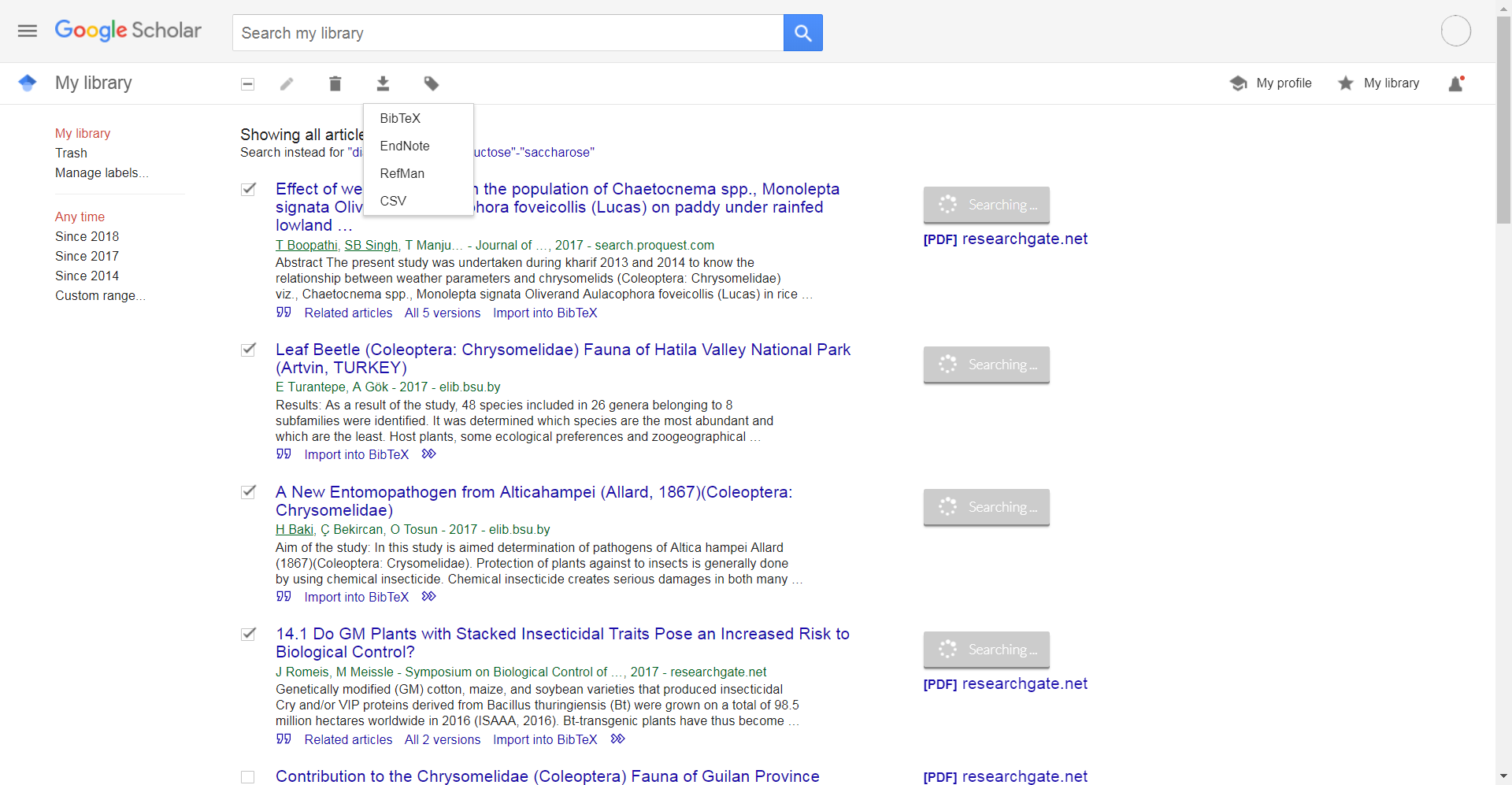

Figure 10.

8. Emptying cache. **DO NOT do this before you have run the searches with cache**
   a. Google Chrome
      - Please see <https://www.technipages.com/google-chrome-clear-cache>
   b. Mozilla Firefox
      - <https://support.mozilla.org/en-US/kb/how-clear-firefox-cache>
   c. Internet Explorer
      - <https://kb.wisc.edu/page.php?id=15141>
9. Abbreviations of institutions:

| **Fujian Agriculture and Forestry University** | **FAFU** |
| --- | --- |
| **Aarhus University** | **AU** |
| **Brock University** | **BU** |
| **Charles Sturt University** | **CSU** |
| **Zhejiang University** | **ZU** |
| **Eötvös Loránd University** | **ELTE** |
| **University of Pecs** | **PTE** |
| **NSW Department of Primary Industries** | **NDPI** |
| **Cambridge University** | **CU** |
| **The James Hutton Institute** | **JHI** |
| **University of Goettingen** | **UG** |
| **University of Kentucky** | **UK** |
